# Selective metal ion utilization contributes to the transformation of the activity of yeast polymerase η from DNA polymerization toward RNA polymerization

**DOI:** 10.1101/2020.09.18.303297

**Authors:** Eva Balint, Ildiko Unk

## Abstract

Polymerase eta (Polη) is a translesion synthesis DNA polymerase directly linked to cancer development. It can bypass several DNA lesions thereby rescuing DNA damage-stalled replication complexes. We previously presented evidence implicating *Saccharomyces cerevisiae* Polη in transcription elongation, and identified its specific RNA extension and translesion RNA synthetic activities. However, RNA synthesis by Polη proved rather inefficient under conditions optimal for DNA synthesis. Searching for factors that could enhance its RNA synthetic activity, we have identified the divalent cation of manganese. Here we show, that manganese triggers drastic changes in the activity of Polη. It increases the efficiency of ribonucleoside incorporation into RNA by ∼400-2000-fold opposite undamaged DNA, and ∼3000 and ∼6000-fold opposite TT dimer and 8oxoG, respectively. Importantly, preference for the correct base is maintained with manganese during RNA synthesis. In contrast, activity is strongly impaired, and base discrimination almost lost during DNA synthesis by Polη with manganese. Moreover, Polη shows strong preference for manganese during RNA synthesis even at 25-fold excess magnesium concentration. Based on these, we suggest that selective metal cofactor usage plays an important role in determining the specificity of Polη during synthesis enabling it to function at both replication and transcription.

## INTRODUCTION

DNA polymerases possess catalytic activity to synthesize DNA in a template dependent fashion using deoxy-ribonucleoside-triphosphates (dNTPs). However, the attributes of their activities differ considerably reflecting their diverse cellular functions. Replicative DNA polymerases are responsible for faithful duplication of the genome and because of that they have a highly selective and restrictive active center ensuring that the correct complementary deoxy-ribonucleoside-monophosphates (dNMPs) are inserted into the growing DNA strand. Therefore, it was of surprise that high fidelity replicative DNA polymerases could incorporate ribonucleoside-monophosphates (rNMPs) with relatively high frequency into DNA due to incomplete exclusion of ribonucleoside-triphosphates (rNTPs) from their active centers (1). Though rNMP incorporation occurs with much reduced efficiency compared to dNMP, it has been estimated that replicative DNA polymerases incorporate ∼10,000 rNMPs into the genome of a yeast cell during a single round of replication, putting rNMPs among the most abundant DNA lesions. The excess presence of rNMPs in DNA went undetected for a long time because they are efficiently removed by ribonucleotide excision repair (2)(3). Beside replicative polymerases, several other DNA polymerases have been shown to be able to utilize rNMPs during synthesis (4)(5)(6)(7).

Due to their high selectivity, modifications or lesions in the template strand hinder the movement of replicative DNA polymerases during replication, leading to cell death if unattended. However, translesion synthesis (TLS) DNA polymerases evolved that are capable of synthesizing through DNA lesions. These polymerases can take over synthesis from stalled replicative DNA polymerases and carry out synthesis across lesion sites this way maintaining the continuity of replication (8). Contrary to replicative DNA polymerases, the active centers of TLS DNA polymerases are more spacious and less selective enabling them to accommodate damaged, modified nucleotides. As a result of that, TLS DNA polymerases are error-prone on undamaged DNA frequently inserting non-complementary nucleosides, which can lead to mutagenesis. Therefore, strict regulatory mechanisms restricting their activities to DNA damage sites are visualized. DNA polymerase η (Polη) is a TLS DNA polymerase that is uniquely able to carry out efficient and error-free bypass of the most frequent ultraviolet (UV) light-induced DNA lesions, cyclobutane pyrimidine dimers (9). The importance of this activity is well emphasized by the fact that inactivity of Polη in humans causes the cancer-prone *xeroderma pigmentosum variant* (XP-V) disorder (10)(11). Polη carries out mostly error-free bypass of one of the most frequent spontaneous oxidative DNA lesions 8-oxoguanine (8-oxoG) as well, and it was shown to bypass several other DNA lesions with varying fidelity (12)(8). We and others have recently identified the ability of Polη to use rNTPs during synthesis (13)(14)(15)(16). Like other DNA polymerases, Polη incorporates rNMPs during DNA synthesis with a very low efficiency. Even so, experiments with both human and yeast Polη showed that they could contribute to the accumulation of ribonucleosides in the genomes of human and yeast cells (17)(18). Unexpectedly, our results indicated that the RNA synthetic activity of yeast Polη was specific as it inserted rNMPs at least 10-fold more efficiently into RNA over DNA (13). During RNA extension it could even perform TLS opposite a TT-dimer and 8-oxoG in an error-free manner. Moreover, we found that the lack of Polη impaired transcription elongation and caused transcriptional inhibition of several genes. These findings suggested a role for Polη during transcription elongation, and possibly in TLS during transcription. However, we also found that Polη utilized dNTPs with much higher efficiency than rNTPs during RNA extension. Even though in yeast cells rNTP concentrations are in the millimolar, whereas dNTP concentrations are in the micromolar range, Polη would still synthesize a mixed strand consisting of ribo-and deoxyribonucleosides, which would have detrimental effects on cells. Hence, we surmised that certain cellular factors could improve the specific RNA extension activity of Polη.

DNA polymerases apply a mechanism based on two divalent metal cations during catalysis (19). The catalytic and nucleotide metal-binding sites at their active center coordinate two metal ions that facilitate the nucleophilic attack by the 3’-OH group of the primer on the β-phosphate of the incoming nucleotide. The presence of a third metal ion at the active center has been recently discovered that is probably needed to reduce the product release barrier (20)(21)(22). Mg^2+^ is presumed to be the physiological metal cofactor for DNA polymerases due to its widespread occurrence in nature, its much higher cellular concentration compared with other divalent metal cations, and that *in vitro* it universally activates DNA polymerases. However, other metal ions like Mn^2+^, Co^2+^, and Ni^2+^ can substitute for Mg^2+^ in *in vitro* polymerization reactions, but the replacement usually significantly diminishes either the efficiency or the fidelity of the enzyme, or both (23)(24). Notwithstanding, Polβ, Polβ, Polµ, Polβ, and PrimPol represent the growing group of exemptions where the activity is improved with the replacement metal. For example, human Polβ exhibited higher reactivity in the presence of Mn^2+^ as compared with Mg^2+^ so that it could even extended a blunt-ended double-stranded DNA template (25). Kinetic and thermodynamic analysis suggested that Polβ evolved as a Mn^2+^-specific enzyme (26). Low concentration of Mn^2+^ had positive effects on both the efficiency and the fidelity of gap-filling synthesis by Polµ using either dNTPs or rNTPs (27). Similarly, at low Mn^2+^ concentrations Polβ was more active with either dNTPs or rNTPs (28). PrimPol was also found to be a Mn^2+^-dependent enzyme as its DNA primase and polymerase activities, as well its DNA primer/template binding affinity significantly improved upon Mn^2+^-binding (29).

In the present study we report, that substitution of manganese for the metal cofactor magnesium transforms the activity of Polη. It greatly impairs its activity and sharply decreases the fidelity of the enzyme during DNA synthesis. In contrast, RNA synthesis becomes 400-2000 fold more efficient with manganese concomitantly maintaining the base selectivity of the enzyme. Moreover, the weak damage bypass activity of Polη observed during RNA synthesis with magnesium is augmented by 3000-6000-fold with manganese opposite TT dimer and 8-oxoG, respectively. Also of note, manganese is preferred by Polη over magnesium even at 25-fold lower concentration during RNA synthesis. Based on these findings we propose a model with a new regulatory mechanism contributing to the shift between the DNA and RNA synthetic activities of Polη.

## MATERIALS AND METHODS

### Protein purification

*Saccharomyces cerevisiae* Polη was overexpressed in yeast in N-terminal fusion with GST and affinity purified on glutathione–Sepharose beads as described previously (13). The GST-tag was removed in the last step of the purification by incubating the beads with PreScission protease. Efficiency of the purification was verified by polyacrylamide gel electrophoresis and Coomassie staining.

### Oligonucleotides and primer extension assays

Sequences of DNA/DNA and RNA/DNA primer/template substrates used in this study are shown in Table S1. Oligonucleotides used as primers contained a fluorophore indocarbocyanine (Cy3) label at the 5’-ends. Oligonucleotides used in these experiments were purchased from Integrated DNA Technologies, Coralville, Iowa, except for the 8-oxoG containing primer that was from Midland Certified Reagent Co. Midland, Texas, and the TT-dimer containing oligonucleotide was from Trilink Biotechnologies, San Diego, California. Results were also verified with DNA and RNA primers purchased from Sigma-Aldrich Merck KGaA, Darmstadt, Germany. Standard primer extension reactions (5 μl) contained 25 mM Tris/HCl pH 7.5, 1 mM dithiothreitol, 100 μg/ml bovine serum albumin, 10% glycerol, the specified divalent cation as chloride salt, and substrate and enzyme as described in the figure legends. Reactions were initiated by the addition of the cation at the indicated concentrations, incubated at 30 °C and quenched by the addition of 15 μl loading buffer containing 95% formamide, 18 mM EDTA, 0.025% SDS, 0.025% bromophenol blue and 0.025% xylene cyanol. The reaction products were resolved on 10-14 % polyacrylamide gels containing 7 M urea and analyzed with a Typhoon TRIO Phosphorimager (GE Healthcare).

### Determination of steady-state kinetic parameters

Primer extension reactions were performed as described above with the following modifications. On undamaged templates, 1 nM Polη was incubated with 20 nM of primer substrate in standard buffer containing 5 mM Mn. Reactions were initiated by adding the corresponding single rNTP (varied from 0.05 to 500 µM) and incubated at 30 °C from 30 sec to 2 min. For kinetic analysis of 8-oxoG or TT dimer bypass, 1 nM Polη was incubated with 8 nM or 16 nM primer substrate, respectively, in standard buffer containing 5 mM Mn. Reactions were initiated by adding rCTP (0.05 to 500 µM) or rATP (0.05 to 500 µM), and incubated at 30 °C for 3 and 10 min, respectively. The intensity of the gel bands corresponding to the substrate and the product were quantitated with Typhoon TRIO Phosphorimager (GE Healthcare) using ImageQuant TL software (GE Healthcare) and the observed rates of nucleotide incorporation were plotted as a function of rNTP concentration. The data were fit by nonlinear regression using SigmaPlot program (version 12.5 Systat Software, San Jose, CA) to the Michaelis-Menten equation describing a hyperbola, *v* = *V*max*[rNTP]/(*Km* + [rNTP]). The *k*_*cat*_ and *K*_*m*_ steady-state parameters were obtained from the fit and were used to calculate the efficiency (k_cat_/K_m_) and the relative efficiency (activation by Mn^2+^ versus Mg^2+^) using the formula f_rel_ = (k_cat_/K_m_)_Mn2+_/(k_cat_/K_m_)_Mg2+_.

## RESULTS

### Effect of divalent metal ions on the synthetic activity of Polη

In our attempt to unravel conditions that could enhance the RNA synthetic activity of Polη, we tested the metal ion dependence of both of its DNA and RNA synthetic activities. Besides magnesium, we compared its activities in the presence of 6 other divalent metal cations that had been implicated in enzymatic activation. In *in vitro* primer extension assays each tested cation supported dNMP insertion into both DNA and RNA primers to a varying extent, although Polη exhibited the highest activity in the presence of Mg^2+^ and Mn^2+^ lengthening almost all the primers, and synthesizing till the end of the template (Fig. 1B and D). In case of each metal cofactors, activation could be detected even at low 0.5 mM metal concentration, whereas high 5 mM metal concentration resulted in higher activity. Surprisingly, when the reactions were supplemented with rNTPs instead of dNTPs, activity was observed only with Mn^2+^ at 0.5 mM metal ion concentration using either a DNA or an RNA primer (Fig. 1C and E). Moreover, Polη exhibited strong activity only in the presence of Mn^2+^ even at 5 mM metal concentration, and only a weak activity could be detected with Mg^2+^, Fe^2+^, and Co^2+^, in agreement with our previous results, whereas no activity was detected with the other metals, Ca^2+^, Ni^2+^, and Zn^2+^ (13). These data suggested that Mn^2+^ could be the proper cation needed for the activation of the RNA synthetic activity of Polη.

**Figure 1.**
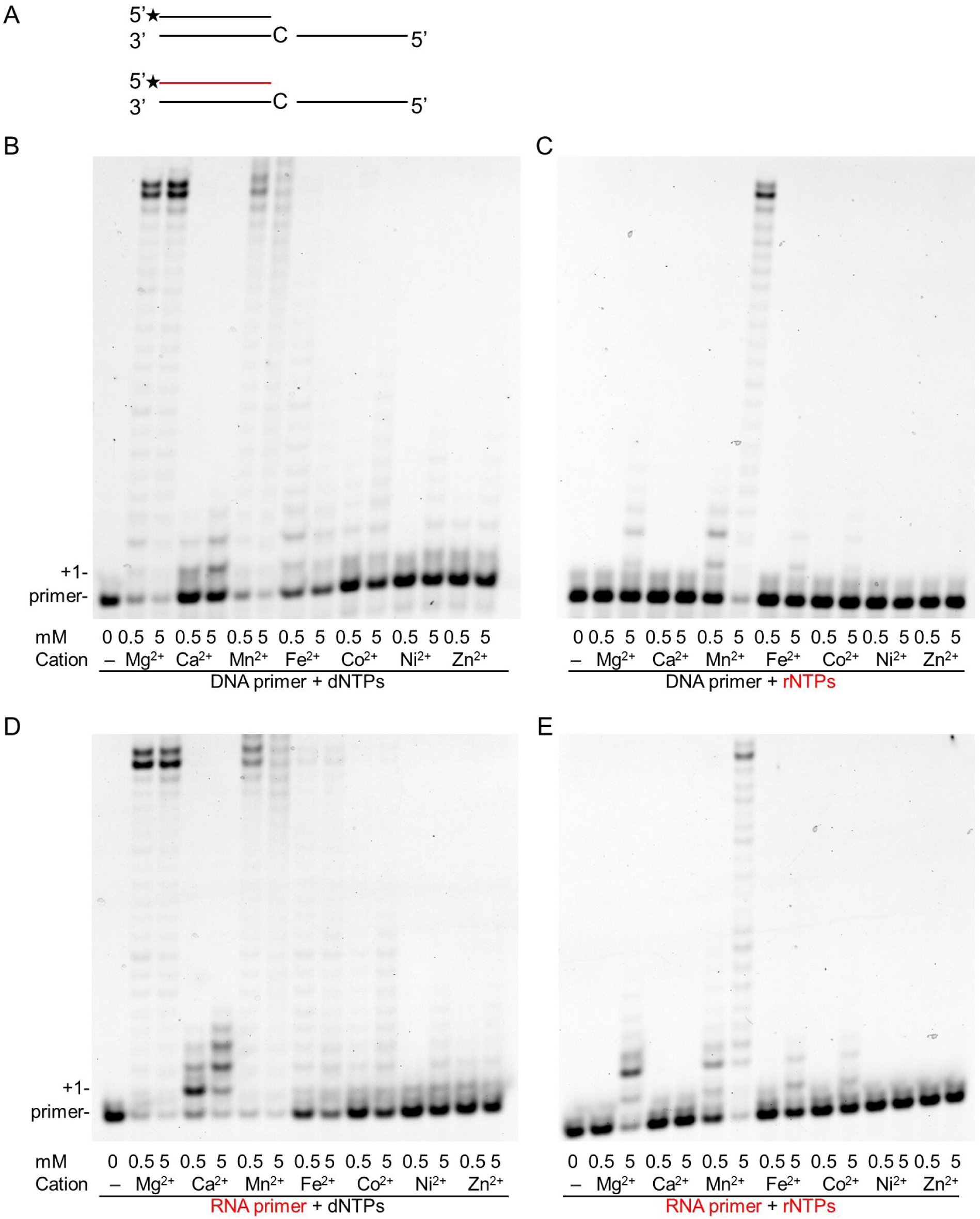
Effect of various divalent cations on the primer extension activities of Polη. *In vitro* primer extension reactions were performed with 40 nM Polη for 5 minutes in the presence of two different concentrations of the indicated divalent cations. (A) The structures of the primer/template used in the experiments are shown. The RNA primer is depicted in red. Asterisks (*) indicate fluorescently labeled primer ends. Reactions were carried out in the presence of DNA primer (S4) and (B) dNTPs or (C) rNTPs, or RNA primer (S8) with (D) dNTPs or (E) rNTPs. Reactions in (B-E) contained 30 nM of the hybridized primer/template, and close to physiological concentrations of nucleotides, either 50 µM dNTPs, or 1 mM rNTPs. The positions of the primer and its extension by one nucleotide are indicated. RNA primers and rNTPs are highlighted in red.

### Mg^2+^ and Mn^2+^ concentration dependent synthesis by Polη

To compare the effect of Mg^2+^ and Mn^2+^ on the synthetic activities of Polη, we applied increasing concentrations of the metal ions and carried out synthesis reactions using a DNA or an RNA primer with dNTPs or rNTPs, in all four combinations. In these experiments, the highest synthetic activities were observed at the highest applied (5 mM) metal ion concentrations in all primer-substrate combinations with both Mg^2+^ and Mn^2+^ (Fig. 2). As expected, in the presence of a DNA primer and Mg^2+^ Polη exhibited high activity with dNTPs, which sharply increased with increasing Mg^2+^ concentrations (Fig. 2A, lanes 6-9). At the highest 5 mM Mg^2+^ concentration Polη extended ∼90 % of the primers and synthesis reached the end of the template. Interestingly, similarly strong activity was observed when the DNA primer was replaced with an RNA primer (Fig. 2C, lanes 6-9). On the other hand, substituting Mn^2+^ for Mg^2+^ resulted in greatly diminished dNMP insertions by Polη on both DNA and RNA primers (Fig. 2A and C, lanes 1-5). Though higher Mn^2+^ concentration resulted in somewhat higher activity, it did not become considerably stronger even at the highest Mn^2+^ concentration extending only ∼30 % of the primers. Contrary to the low activation of DNA synthesis, Mn^2+^ dramatically enhanced rNMP insertion by Polη utilizing either a DNA or an RNA primer. Whereas rNMP incorporation was inefficient with Mg^2+^ extending ∼10 % of the primers with only 1-2 nucleotides at the highest Mg^2+^ concentration (Fig. 2B and D, lanes 6-9), it sharply increased with increasing Mn^2+^ concentration lengthening ∼90 % of the primers and resulting in fully extended primers at 5 mM Mn^2+^ concentration (Fig. 2B and D, lanes 1-5). The same Mn^2+^ concentration-dependent activation could be detected using individual rNTPs in the assays (Fig. 2E and F, and Fig. S1). In summary, these results showed that although Mn^2+^ strongly reduced the DNA synthetic activity of Polη compared to Mg^2+^, it dramatically elevated its RNA synthetic activity in a concentration dependent manner.

**Figure 2.**
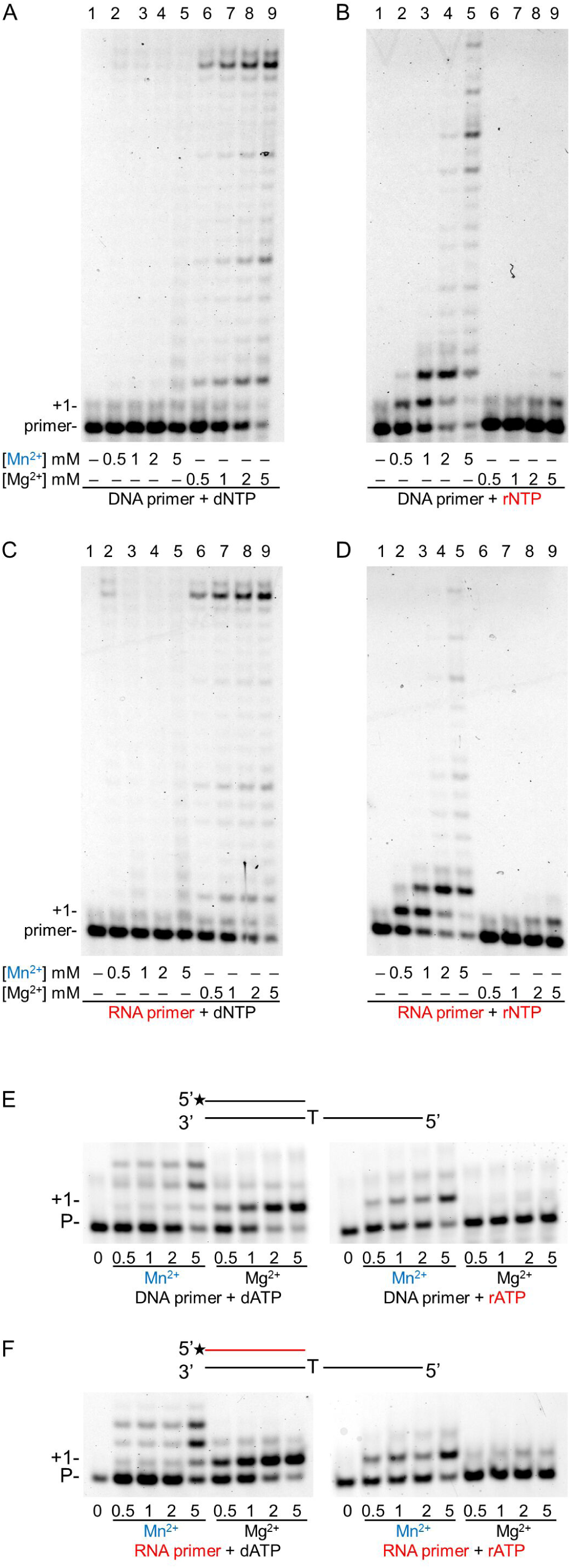
The effect of increasing concentrations of magnesium and manganese ions on the primer extension activity of Polη in the presence of dNTPs or rNTPs. (A-D) Reactions were performed with 10 nM Polη in the presence of 20 nM primer/template (S3 for A, B, and S7 for C, D) and close to physiological concentrations of nucleotides, either 50 µ M dNTPs, or 1 mM rNTPs. (A) DNA primer and dNTPs (B) DNA primer with rNTPs (C) RNA primer with dNTPs (D) RNA primer and rNTPs. The concentrations of Mn^2+^ and Mg^2+^ are indicated below each panel. Lanes are numbered at the top. The single ribonucleotide incorporation assays shown on (E-F) contained the indicated concentrations of Mn^2+^ or Mg^2+^, 6 nM Pol η and 50 μM dATP or 1 mM rATP, with (E) 20 nM DNA primer (S2) or (F) 20 nM RNA primer (S6). For (A-F) reactions were incubated for 5 minutes. The positions of the primer and its extension by one nucleotide are indicated. RNA primers and rNTPs are highlighted in red, and manganese in blue.

In order to determine the proper concentration of Mn^2+^ needed for the highest activation of RNA synthesis, we tested reactions containing Mn^2+^ in the broad range of 0.1-25 mM. As Fig. 3 shows, Polη was activated at all Mn^2+^ concentrations, and the highest activity was detected with 5 mM Mn^2+^. Lower or higher concentrations resulted in gradually decreasing activity, though the changes were more moderate at the higher range.

**Figure 3.**
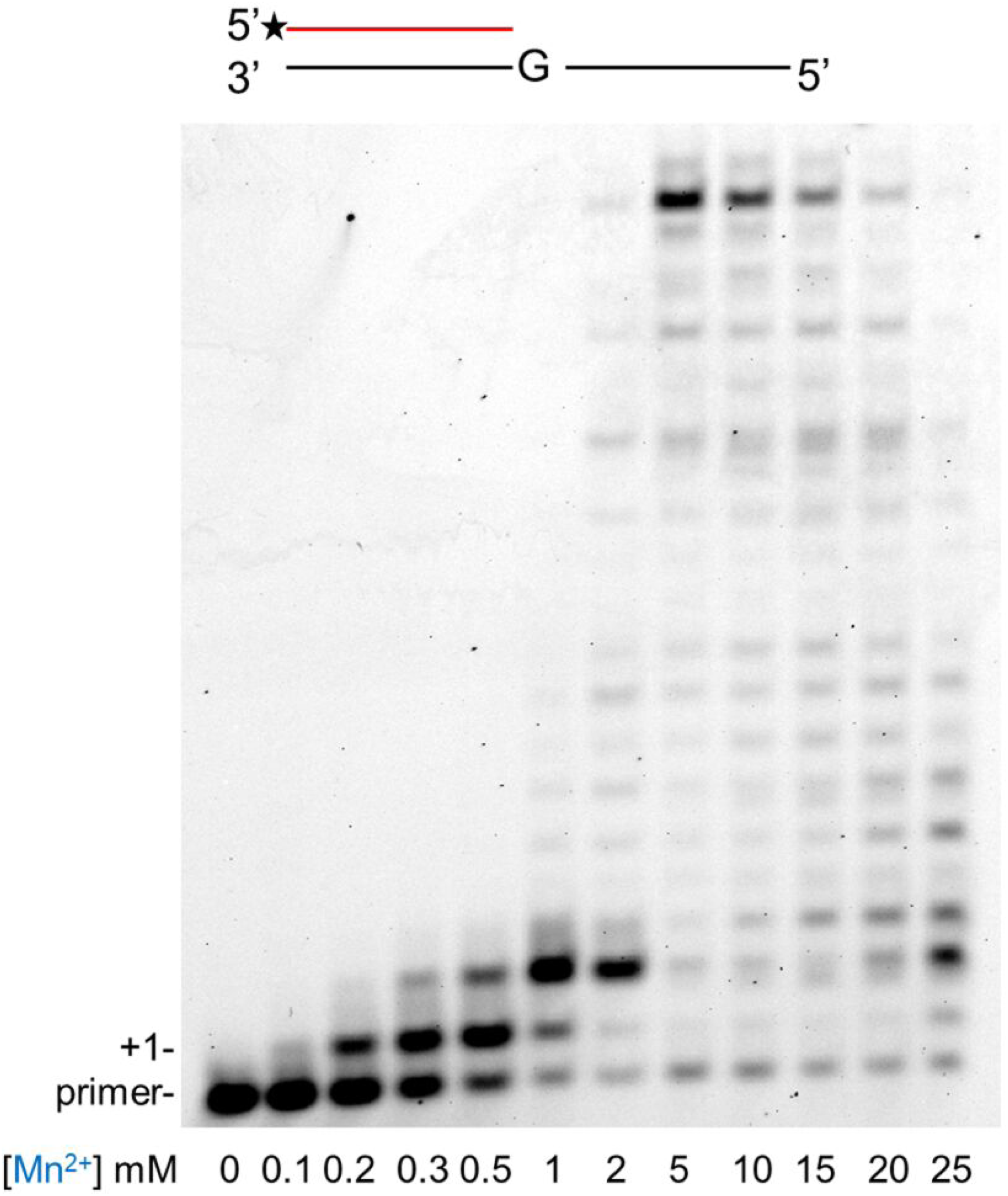
Optimal manganese concentration needed for the highest activation of Polη during RNA synthesis. Primer extension reactions using S8, containing the indicated concentration of Mn^2+^ were performed for 5 minutes with 20 nM Pol η and 1 mM rNTP. The positions of the primer (P) and its one nucleotide extention (+1) are indicated.

### Kinetics of correct rNMP incorporation into RNA in the presence of Mn^2+^

Previously, we determined the kinetic parameters of the RNA synthetic activity of Polη in the presence of Mg^2+^ and found that it incorporated single rNMPs into RNA with an efficiency of ∼10^−3^-10^−4^ min^−1^µM^−1^ (13). To quantitate the enhancement of its RNA synthetic activity observed in the presence of Mn^2+^, we carried out similar steady-state kinetic studies using 5 mM Mn^2+^ instead of 5 mM Mg^2+^ in the reactions. Remarkably, as Table 1 shows, Polη inserted rNMPs into RNA ∼1000 fold more efficiently when utilizing Mn^2+^. The smallest ∼400-fold increase was detected during rCMP insertion, whereas the highest ∼2000-fold difference was measured during rAMP insertion. The overall enhancement was due to a ∼100-fold decrease in the apparent K_m_ indicating stronger rNTP-binding of Polη, and to a ∼10-fold increase in the k_cat_ values reflecting the velocity of the reactions. These observed changes in the K_m_ and k_cat_ values indicated a greatly improved specificity of the RNA extension reactions achieved in the presence of Mn^2+^.

**Table 1.**
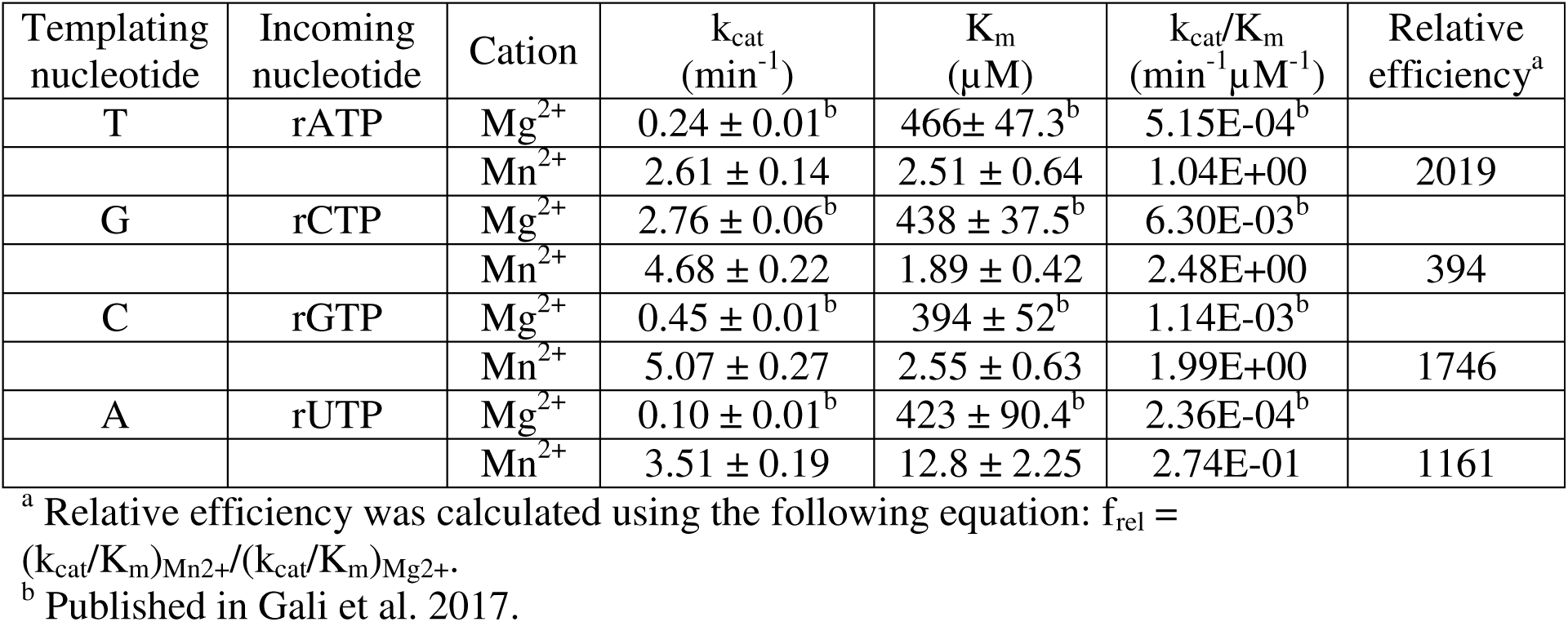
Comparison of the kinetic parameters of rNTP incorporation into RNA by Polη using Mg^2+^ or Mn^2+^ as cofactor.

### The effect of Mn^2+^ on the base selectivity of Polη

*In vitro* Mn^2+^ can substitute for Mg^2+^ in the activation of several DNA polymerases. However, in most cases the accuracy of DNA synthesis supported by Mn^2+^ drastically decreases spoiling the activity of the polymerase. Hence, we investigated the effects of Mn^2+^ on the base selectivity of Polη by testing the incorporation of all 4 dNMPs and rNMPs individually into DNA and RNA primers, respectively, opposite each 4 possible DNA template residues. When DNA synthesis was assayed using Mg^2+^ Polη showed preference for the correct dNTP in accordance with its reported 10^−2^-10^−4^ fidelity (Fig. 4A). However, base selectivity was almost completely lost with Mn^2+^ and the correct and incorrect dNMPs were inserted with comparably weak efficiencies (Fig. 4A). During RNA synthesis with Mg^2+^ Polη discriminated against the incorrect bases catalyzing only weak misinsertions as opposed to robust correct rNMP insertions (Fig. 4B). Surprisingly, although Mn^2+^ resulted in somewhat decreased base selectivity indicated by the stronger intensity of the bands representing misinsertions, still clear preference for the correct rNTPs was maintained (Fig. 4B). To reconfirm these data, we quantitated the fidelity of RNA synthesis in the presence of Mn^2+^ in steady-state kinetic experiments (Fig. S2). The results of these studies showed that Polη exhibited base selectivity in the 10^−2^-10^−4^ range during RNA synthesis with Mn^2+^, which corresponded well with its reported accuracy during DNA synthesis with Mg^2+^ (Table 2) (30). Interestingly, rAMP misincorporations were the weakest (∼10^−3^-10^−4^), whereas rCMP was misinserted with the highest (10^−1^-10^−2^) relative efficiencies opposite all three non-complementary template residues. In summary, the above data indicated that Polη maintained its fidelity during RNA synthesis in the presence of Mn^2+^.

**Table 2.** Kinetic parameters of rNTP incorporation and misincorporation into RNA by Polη using Mn^2+^ as cofactor.

| Templating nucleotide | Incoming nucleotide | k<sub>cat</sub> (min<sup>-1</sup>) | K<sub>m</sub> (μM) | k<sub>cat</sub>/K<sub>m</sub> (min<sup>-1</sup>μM<sup>-1</sup>) | Relative efficiency<sup>a</sup> | Discrimination 1/f<sup>b</sup> |
| --- | --- | --- | --- | --- | --- | --- |
| T | rATP | 2.61± 0.14 | 2.51± 0.64 | 1.04 |  |  |
|  | rCTP | 1.72± 0.06 | 19.4± 3.14 | 0.09 | 8.7E-02 | 12 |
|  | rGTP | 0.72± 0.03 | 44.6 ± 7.03 | 0.016 | 1.5E-02 | 67 |
|  | rUTP | 0.36± 0.02 | 261 ± 50.6 | 0.0014 | 1.3E-03 | 769 |
| G | rATP | 0.06± 0.00 | 89.3± 17.7 | 0.0007 | 2.8E-04 | 3571 |
|  | rCTP | 4.68± 0.22 | 1.89± 0.42 | 2.48 |  |  |
|  | rGTP | 0.31± 0.02 | 68.9 ± 14.9 | 0.0045 | 1.8E-03 | 555 |
|  | rUTP | 1.86± 0.04 | 199 ± 14.5 | 0.0093 | 3.8E-03 | 263 |
| C | rATP | 0.10± 0.00 | 69.2± 11.9 | 0.0014 | 7.0E-04 | 1429 |
|  | rCTP | 1.03± 0.04 | 19.9± 3.74 | 0.052 | 2.6E-02 | 38 |
|  | rGTP | 5.07± 0.27 | 2.55± 0.63 | 1.99 |  |  |
|  | rUTP | 0.16± 0.01 | 339 ± 49.4 | 0.0005 | 2.5E-04 | 4000 |
| A | rATP | 0.05± 0.00 | 138± 26.2 | 0.0004 | 1.48E-03 | 676 |
|  | rCTP | 1.03± 0.07 | 17.5± 4.92 | 0.059 | 2.2E-01 | 5 |
|  | rGTP | 0.24± 0.01 | 55.9± 8.64 | 0.0043 | 1.6E-02 | 63 |
|  | rUTP | 3.51± 0.19 | 12.8± 2.25 | 0.27 |  |  |

<sup>a</sup> Relative efficiency was calculated using the following equation: $f_{rel} =$
<sup>b</sup> Inverse relative efficiency: $1/f_{rel} = (k_{cat}/K_m)_{correct}/(k_{cat}/K_m)_{incorrect}$

**Figure 4.**
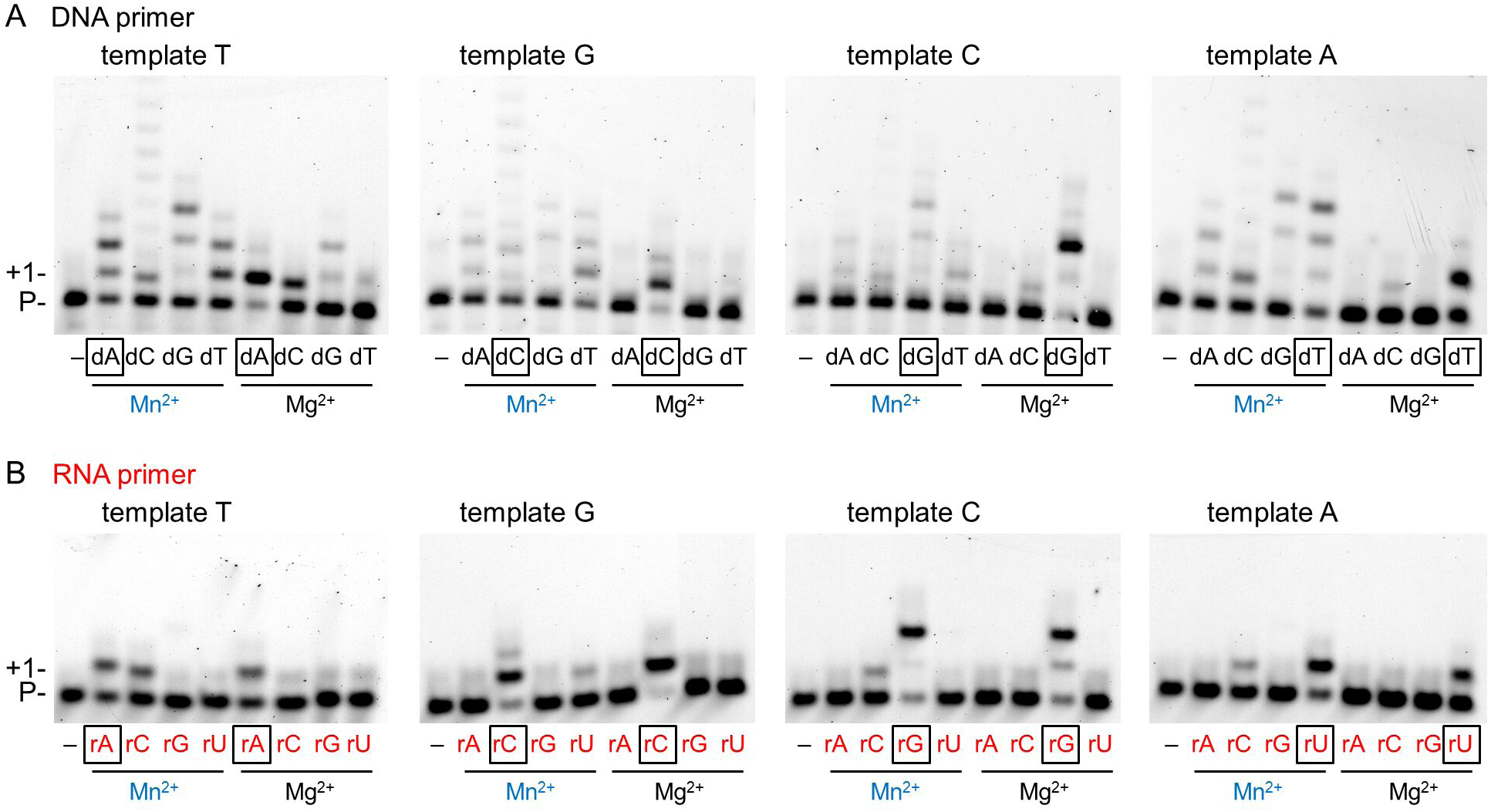
Fidelity of Pol η during DNA and RNA synthesis in the presence of magnesium or manganese. Reactions contained 5 mM Mn^2+^ or Mg^2+^ as indicated, 6 nM Pol η, 20 nM (A) DNA/DNA (S1-4) or (B) RNA/DNA (S5-8) primer/template, and (A) 0.1 mM individual dNTPs or (B) 4 mM individual rNTP, as indicated. Reaction times were 1 min except for the 15 min reactions with Mg^2+^ on (B). The templating base in the incoming position is denoted. RNA primer and rNTPs are highlighted in red, and manganese is in blue. The nucleotides representing correct insertions are boxed.

### DNA damage bypass activity of Polη with Mn^2+^

Next, we examined the effect of Mn^2+^ on the TLS activity of Polη during RNA extension (Fig. 5A and C). Our previous results obtained in the presence of Mg^2+^ revealed a very inefficient bypass of 8-oxoG and TT dimer during RNA extension (13). Importantly, Mn^2+^ had a profound effect on insertion opposite both these damages (Fig. S3). Table 3 shows that a ∼6000-fold and a ∼3000-fold enhancement in TLS efficiency was measured in steady state kinetic experiments opposite 8-oxoG and TT dimer, respectively, compared to data obtained in the presence of Mg^2+^. As in the case of undamaged templates and correct incoming nucleotides, the apparent K_m_ values decreased by 2 orders of magnitude, whereas the K_cat_ values increased by an order of magnitude with Mn^2+^. Moreover, Polη kept its fidelity during the bypass reactions as it preferably inserted the correct rNMPs opposite the damage sites and no significant insertions of the incorrect nucleotides were observed (Fig. 5B and D). Based on these results we concluded that Mn^2+^ was a specific activator of the RNA synthetic activity of Polη both on undamaged templates, and opposite DNA damages.

**Table 3.**
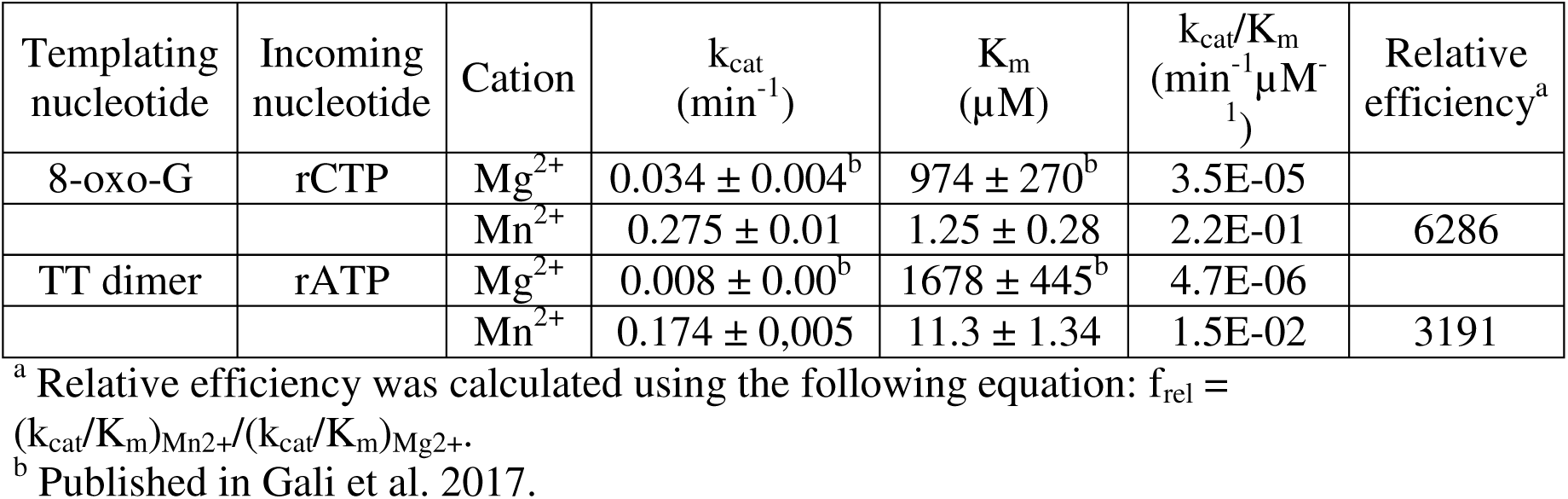
Kinetic parameters of rNTP incorporation into RNA by Polη opposite DNA damages using Mg^2+^ or Mn^2+^ as cofactor.

**Figure 5.**
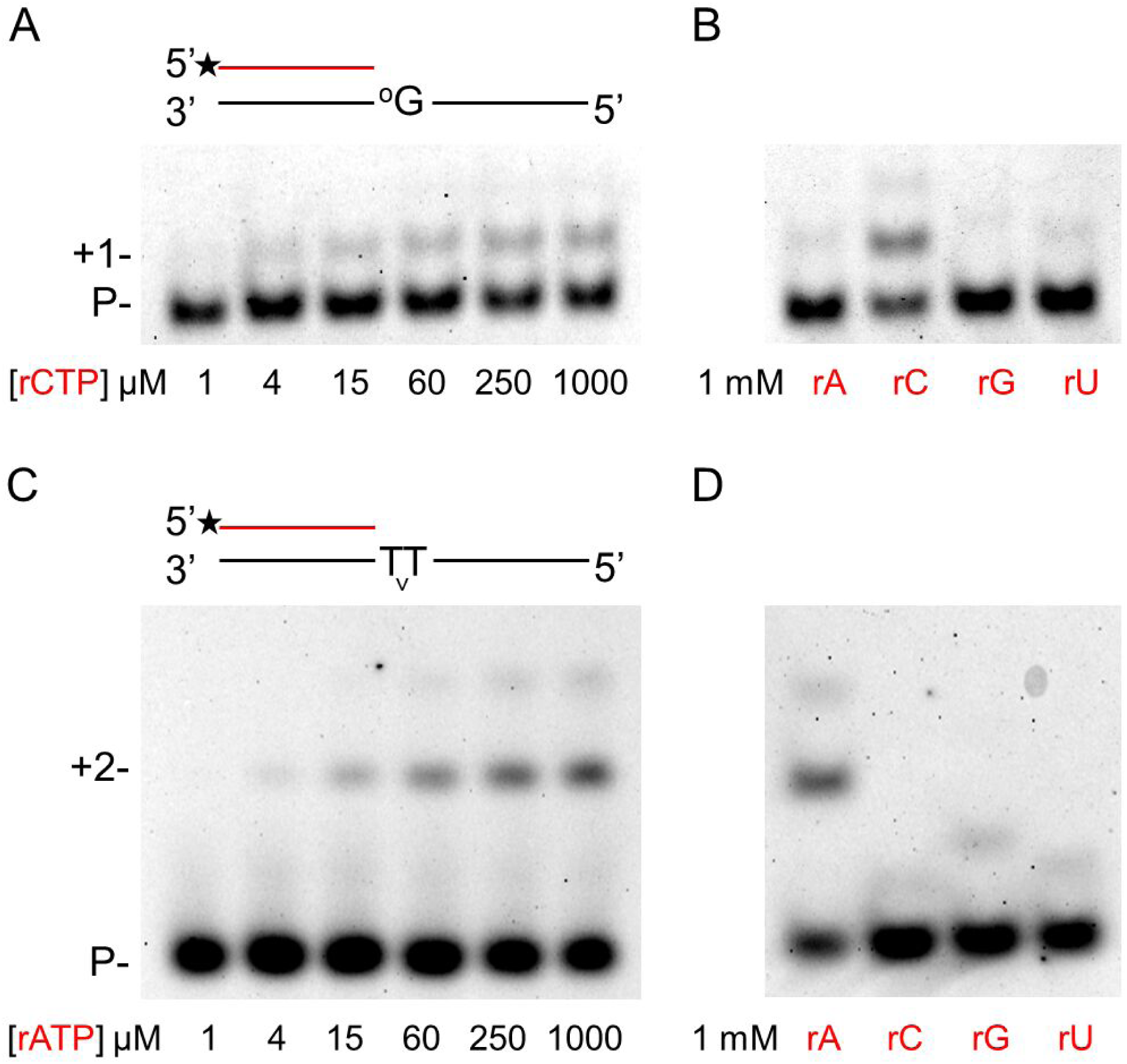
DNA damage bypass by Polη during RNA synthesis in the presence of manganese. (A-B) The template in S12 contained 8-oxoG in the incoming position. Reactions were performed for 3 min with 5 mM Mn^2+^, 1.6 nM Pol η, 8 nM RNA/DNA primer/template and the indicated amount of individual rNTPs. (C-D) The template in S16 contained TT dimer in the incoming position. Reactions were performed for 15 min with 5 mM Mn^2+^, 1.6 nM Pol η, 20 nM RNA/DNA primer/template and the indicated amount of individual rNTPs. RNA primers and rNTPs are highlighted in red.

### Metal preference of Polη during RNA synthesis

Since Mn^2+^ exerted such a dramatic effect on Polη activity, it was important to examine which metal cation was preferred by Polη during RNA synthesis. For this reason, we carried out RNA extension experiments with rNTPs in the joint presence of Mg^2+^ and Mn^2+^. In the first set of reactions, the concentration of Mg^2+^ was gradually decreased from 6 mM to 0 mM, whereas the concentration of Mn^2+^ was increased from 0 mM to 6 mM in parallel maintaining the total metal cation concentration at 6 mM. The pattern of reaction products contrasted strikingly in the sole presence of Mg^2+^ or Mn^2+^ (Fig. 6A, compare the first and last lanes) enabling easy detection of cation utilization. The results showed, that even at eleven-fold excess of Mg^2+^ reaction products specific for Mn^2+^ appeared (Fig. 6A second lane). In the next set of reactions Mg^2+^ concentration was kept at 5 mM and Mn^2+^ concentration was gradually increased. In this setup, the reaction products showed a Mn^2+^-specific pattern already at 0.2 mM Mn^2+^ concentration despite the 25-fold higher Mg^2+^ level indicating that Polη preferred Mn^2+^ over Mg^2+^ in the reactions (Fig 6B, lane 3).

**Figure 6.**
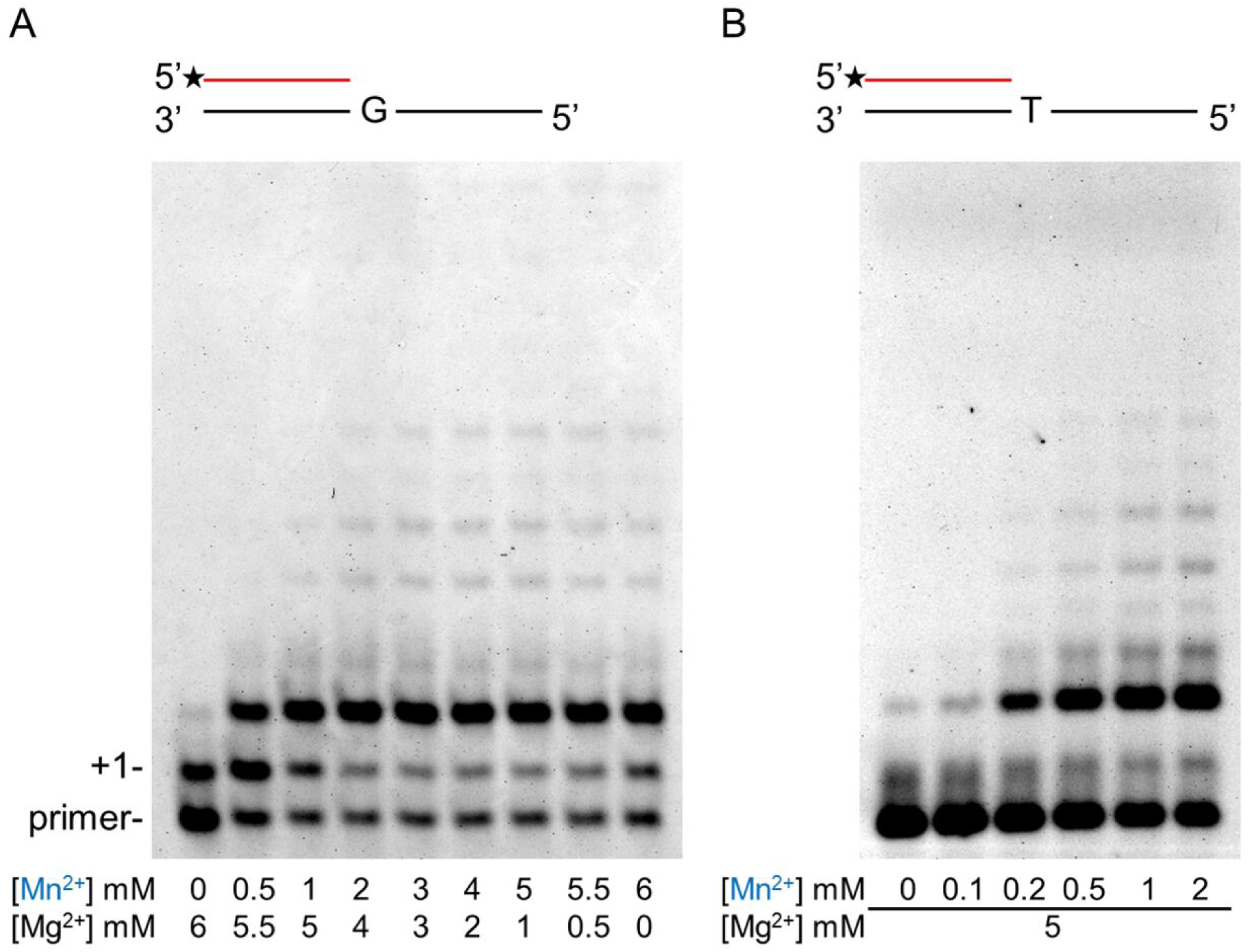
Metal ion preference of Polη during RNA synthesis. Primer extension reactions were performed with 10 nM Pol η, 20 nM RNA/DNA primer/template and 1 mM rNTP mix for 5 min. (A) Reactions were performed with S7 and contained both Mn^2+^ and Mg^2+^ in the indicated concentrations. (B) Reactions using S6 contained 5 mM Mg and the indicated concentrations of Mn^2+^. Manganese is highlighted in blue. The positions of the primer and its one nucleotide extention (+1) are indicated.

## DISCUSSION

The aim of the present study was to identify cellular factors that could improve the RNA synthetic activity of yeast Polη. The conception was based on our previous results showing that Polη has a specific RNA synthetic activity inserting rNMPs at least ten times more efficiently into an RNA primer as opposed to a DNA primer (13). Despite its specificity, the observed efficiency of RNA synthesis was rather weak raising the possibilities that either the applied reaction conditions were not appropriate, or Polη required accessory proteins for efficient RNA synthesis, or both. Hence, first we tried to optimize the reaction by replacing the generally used metal cofactor Mg^2+^ with other divalent metal cations. We tested several metal cations and all could activate DNA synthesis to varying degrees, but Mg^2+^ and Mn^2+^ achieved the highest activity. On the other hand, during RNA synthesis Ca^2+^, Ni^2+^, and Zn^2+^ were inactive, Mg^2+^, Fe^2+^, and Co^2+^ conferred very limited activity, and only Mn^2+^ supported efficient reaction. These results suggest that Mn^2+^ is the proper metal cofactor of Polη during RNA synthesis, and that other metal cations can not substitute for it. Steady-state kinetic experiments revealed that in comparison with Mg^2+^, Mn^2+^ caused a 400-2000-fold increase in efficiency during RNA synthesis on undamaged templates, and a 6000 and 3000-fold increase opposite 8-oxoG and TT dimer, respectively. The specificity of the activation is underpinned by that the enzyme maintained its base selectivity in the 10^−2^-10^−4^ range with Mn^2+^, similarly to its base discrimination during DNA synthesis with Mg^2+^ (30). Moreover, Polη preferentially utilized Mn^2+^ even in a 25-fold excess of Mg^2+^ during RNA synthesis. Taken together these data reinforce our previous finding that the RNA synthetic activity of Polη is specific, and identify Mn^2+^ as its apposite metal cofactor.

Importantly, our experiments also demonstrate that selective utilization of the two metal cations Mg^2+^ and Mn^2+^ results in a complete transformation in specificity. When utilizing Mg^2+^ Polη is proficient in DNA synthesis but very inefficient during RNA synthesis, and preference for the correct base is sustained in both cases. On the other hand, contrary to the remarkable enhancement of RNA synthesis by Mn^2+^, it adversely affected the DNA synthetic activity of Polη by strikingly decreasing both the efficiency and the fidelity of the reaction. This differential effect of Mn^2+^ on the DNA and RNA synthetic activities of Polη sharply contrasts the effect it had on the reported Mn^2+^-dependent polymerases, Pols β, β, µ, and Primpol, in which cases Mn^2+^ exerted an overall positive effect on synthesis with either dNTPs or rNTPs, and Polβ even displayed increased fidelity opposite template T, where with Mg^2+^ it favored G misinsertion (26)(27)(28)(29). The advantage of enhanced dNMP incorporation is obvious during DNA synthesis. Nevertheless, increased rNTP utilization by the above DNA polymerases could also be advantageous to cells. As it was suggested, it would allow faster repair during gap-filling or during the repair of DNA double strand breaks by non-homologous end joining in an environment where rNTPs are more easily accessible because of their high intracellular concentration surpassing dNTP concentration. The incorporated rNMPs could be removed later by ribonucleotide excision repair. During transcription, however, the DNA synthetic activity of Polη has to be repressed and the RNA synthetic activity has to be elevated to avoid excess dNMP insertion into RNA, which could hinder elongation, lead to miscoding, or otherwise could alter key steps of transcription and translation (31)(32)(33)(34)(35)(36). In conclusion, we propose that preferential activation of the DNA or RNA synthetic activity with concomitant impairment of the other through selective metal utilization constitutes a new regulatory mechanism that, with the contribution of other yet unidentified factors, enables Polη to take part in synthesis and DNA damage bypass during replication and also during transcription (Fig. 7). To our knowledge, this is the first study reporting the existence of such a regulatory mechanism in the activation of two separate activities of a eukaryotic polymerase.

**Figure 7.**
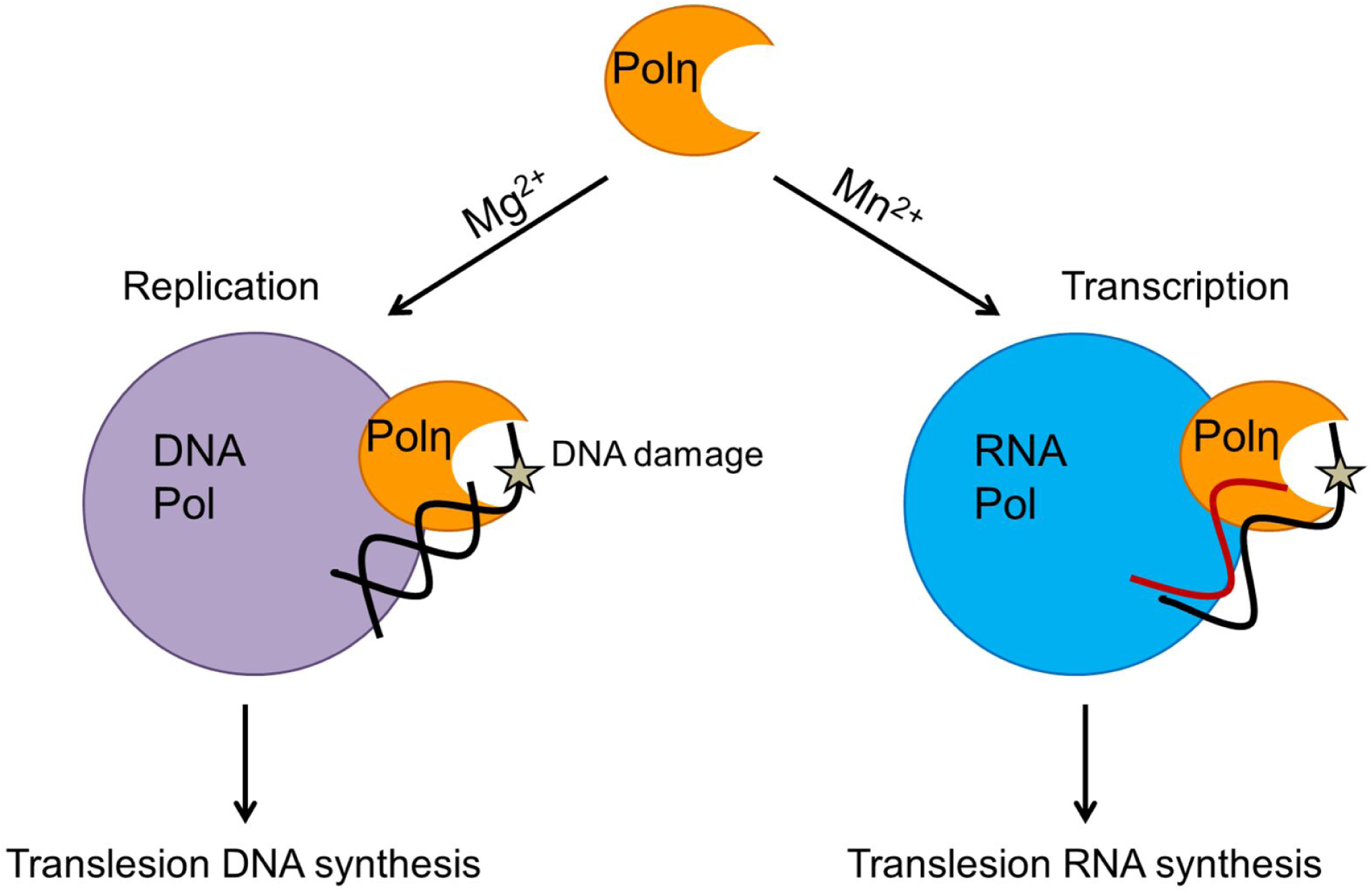
Model for the selective metal cation dependent activities of yeast Polη

Yet, we have to consider that Mg^2+^ is the most abundant divalent metal cation in the cell. The intracellular concentration of Mg^2+^ is in the millimolar range, much higher than the concentration of Mn^2+^ which is in the micromolar range (37). Therefore, Mg^2+^ could be readily acquired by a plethora of enzymes including Polη during DNA replication. In turn, though our results indicate that Mn^2+^ is preferred over Mg^2+^ by Polη during RNA synthesis even at a ∼25-fold lower concentration, given the huge difference between the intracellular concentrations of the two metal ions the involvement of additional factors assisting Mn^2+^-binding has to be presumed. We hypothesize that direct interactions with the transcription machinery could have an effect on the metal utilization of Polη so that Mn^2+^ would be preferred over other cations. Further experiments are needed to unravel the identity of such factor(s) and the way it influences the metal selectivity of yeast Polη.

Yeast and human Polη exhibit very similar biochemical characteristics during DNA synthesis including processivity, fidelity, damage bypass ability. Moreover, human Polη was also shown to be able to utilize rNTPs during DNA extension (15)(16)(17). Therefore, considering its well documented importance in cancer avoidance it would be of high significance to investigate whether human Polη also has Mn^2+^-activated specific RNA synthetic and translesion RNA synthesis activities, adding an additional layer to its contribution to genome stability.

## Acknowledgements

We thank Szilvia Minorits and Aniko Bozo-Toth for technical assistance.

