## Supplemental Table and Figures for "Selective metal ion utilization contributes to the transformation of the activity of yeast polymerase η from DNA polymerization toward RNA polymerization"

**Table S1.**
Sequence and structure of substrates used in the *in vitro* primer extension assays. RNA primers are in red. The Cy3 label at the 5' end of primers is indicated. The first templating nucleotides are in bold.

| Substrate | Sequence |
|---|---|
| S1 | /5Cy3/CGCTACCTAGCCTGCCTCAAGAGTTGCTCG<br>3'-GCGATGGATCGGACGGAGTTCTCAACGAGC**A**CAGGCTTACGCTCAGGTCG-5' |
| S2 | /5Cy3/CGCTACCTAGCCTGCCTCAAGAGTTGCTCG<br>3'-GCGATGGATCGGACGGAGTTCTCAACGAGC**T**CAGGCTTACGCTCAGGTCG-5' |
| S3 | /5Cy3/CGCTACCTAGCCTGCCTCAAGAGTTGCTCG<br>3'-GCGATGGATCGGACGGAGTTCTCAACGAGC**G**CAGGCTTACGCTCAGGTCG-5' |
| S4 | /5Cy3/CGCTACCTAGCCTGCCTCAAGAGTTGCTCG<br>3'-GCGATGGATCGGACGGAGTTCTCAACGAGC**C**CAGGCTTACGCTCAGGTCG-5' |
| S5 | /5Cy3/CGCUACCUAGCCUGCCUCAAGAGUUGCUCG<br>3'-GCGATGGATCGGACGGAGTTCTCAACGAGC**A**CAGGCTTACGCTCAGGTCG-5' |
| S6 | /5Cy3/CGCUACCUAGCCUGCCUCAAGAGUUGCUCG<br>3'-GCGATGGATCGGACGGAGTTCTCAACGAGC**T**CAGGCTTACGCTCAGGTCG-5' |
| S7 | /5Cy3/CGCUACCUAGCCUGCCUCAAGAGUUGCUCG<br>3'-GCGATGGATCGGACGGAGTTCTCAACGAGC**G**CAGGCTTACGCTCAGGTCG-5' |
| S8 | /5Cy3/CGCUACCUAGCCUGCCUCAAGAGUUGCUCG<br>3'-GCGATGGATCGGACGGAGTTCTCAACGAGC**C**CAGGCTTACGCTCAGGTCG-5' |
| S12 | /5Cy3/CGACGAUGCUCCGGUACUCCAGUGUAGGCAU<br>3'-CAAAAGGGTCAGTGCTGCTACGAGGCCATGAGGTCACATCCGTA$^{O}$**G**AATGCTTAAGAA<br>CTCCGTCCGTACCATCGA-5' |
| S16 | /5Cy3/CGUAUUCGCGCGC<br>3'-CGAATGGCGGTGCG**T^T**GCGCGCGAATACG-5' |

**Supplementary Figure legends**

**Supplementary Figure 1.**

Single ribonucleotide incorporation in the presence of manganese or magnesium. Primer extension reactions containing the indicated concentration of $Mn^{2+}$ or $Mg^{2+}$ were performed for 5 minutes with 6 nM Pol η, 20 nM DNA primer/DNA template (S1-4), or RNA primer/DNA template (S5-8), and 50 μM of the incoming individual correct dNTP or 1 mM of the individual correct rNTP, as indicated below the panels. The templating bases are indicated at the top.

**Supplementary Figure 2.**

Determination of the kinetic parameters of rNTP incorporation and misincorporation into RNA primer by Polη using $Mn^{2+}$ as cofactor. Templates contained T (A), G (B), C (C) or A (D) in the incoming position (S5-8), as denoted above the panels. Reactions contained 5 mM $Mn^{2+}$, 20 nM RNA primer/DNA templare, 1 nM Polη (unless otherwise indicated) and increasing concentration of an individual rNTP as shown. Reaction times are also indicated.

**Supplementary Figure 3.**

Determination of the kinetic parameters of DNA damage bypass by Polη using RNA primer and $Mn^{2+}$ as cofactor. A) The template (S12) contained 8-oxoG in the incoming position. Reactions in the presence of 5 mM $Mn^{2+}$, 8 nM RNA primer/DNA templare, 0.8 nM Polη and increasing concentration of rCTP, as shown, were performed for 6 min. B) Reactions with S16 containing a TT dimer in the incoming position were performed for 20 min. in the presence of 5 mM $Mn^{2+}$, 16 nM RNA primer/DNA templare, 1.6 nM Polη and increasing concentration of rATP, as shown.

### Supplementary Figures

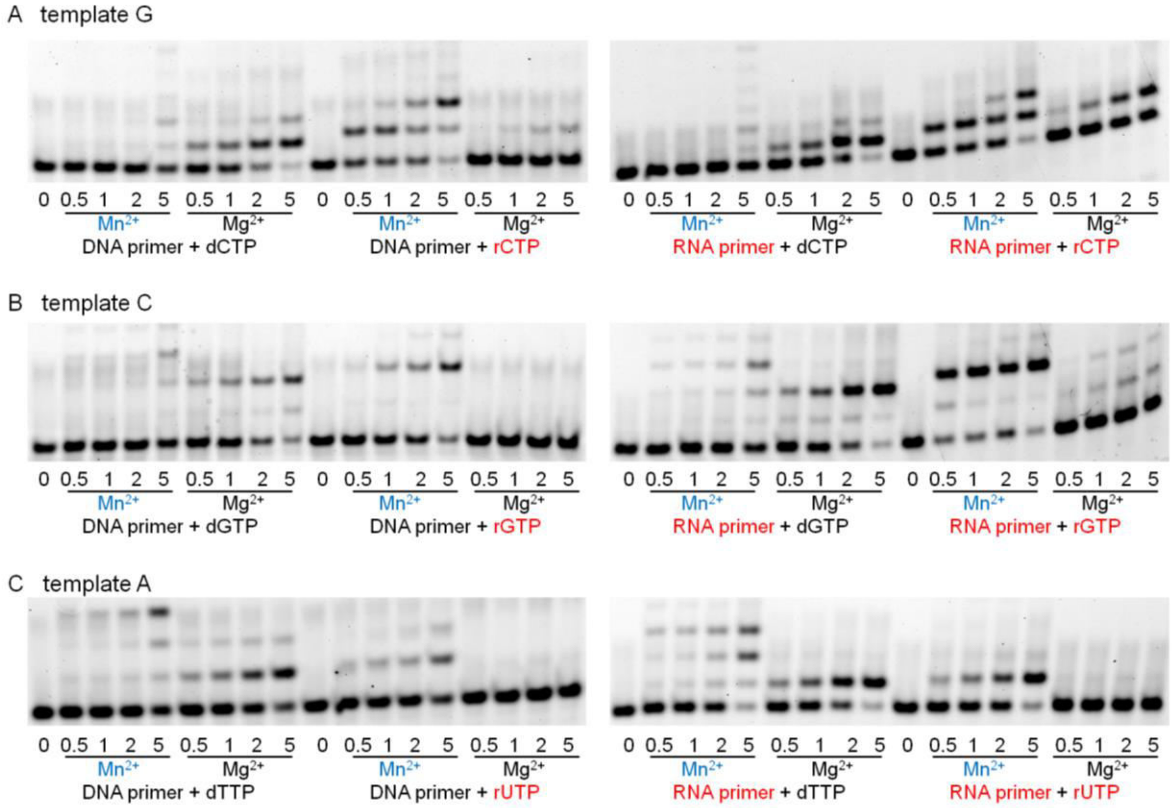


Supplementary Figure 1

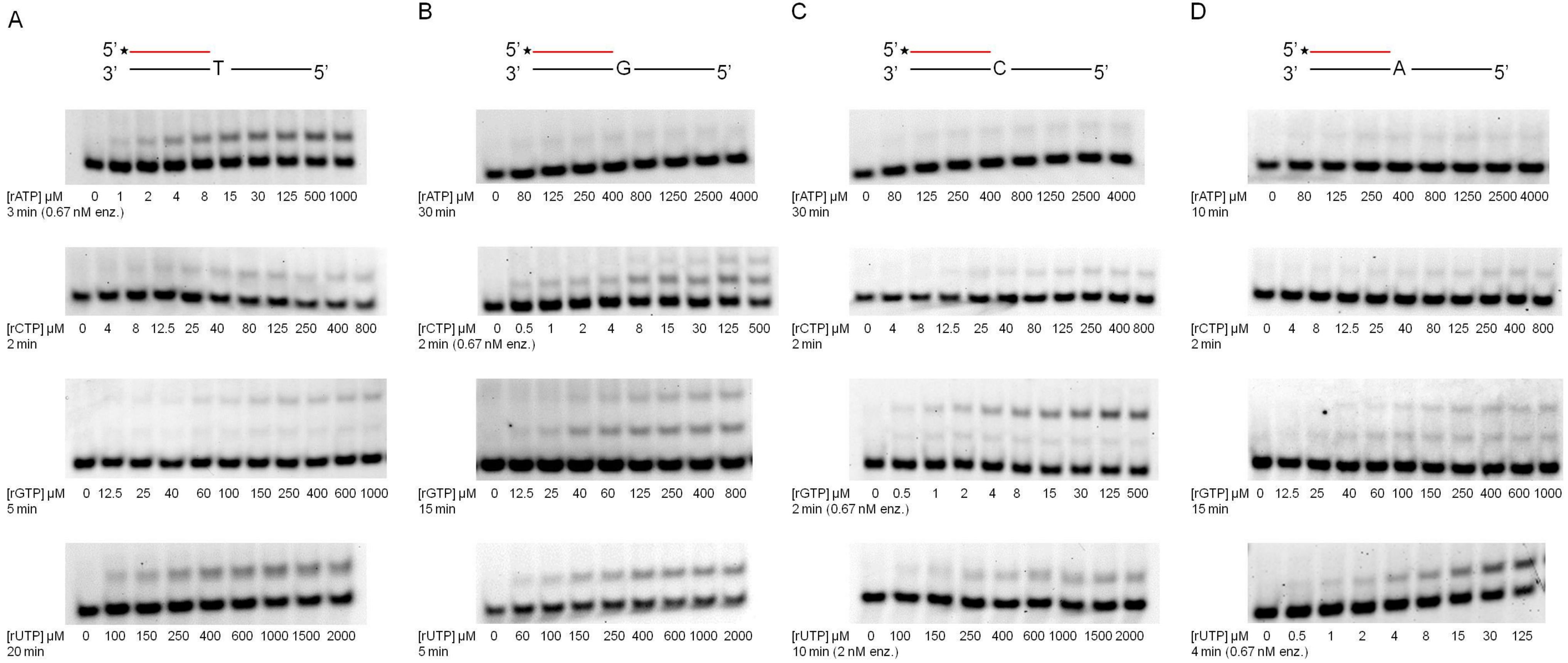


Supplementary Figure 2

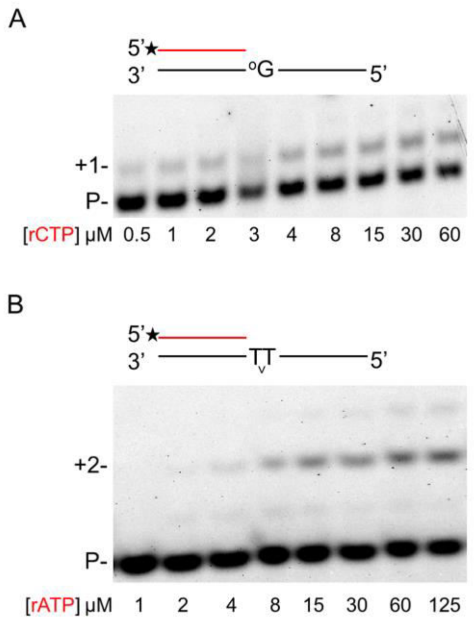


Supplementary Figure 3
